# A Novel Model for Encephalomyosynangiosis Surgery after Middle Cerebral Artery Occlusion-Induced Stroke in Mice

**DOI:** 10.1101/2021.08.16.456489

**Authors:** Mitch Paro, Daylin Gamiotea Turro, Leslie Blumenfeld, Ketan R. Bulsara, Rajkumar Verma

**Affiliations:** Department of Neuroscience, UConn School of Medicine, Farmington, CT; Division of Neurosurgery, UConn School of Medicine, Farmington, CT

## Abstract

**Background and Purpose:** No effective treatment is available for most patients who suffer ischemic stroke. Development of novel treatment options is imperative. The brain attempts to self-heal after ischemic stroke via various mechanism mediated by restored blood circulation in affected region of brain but this process is limited by inadequate angiogenesis or neoangiogenesis. Encephalomyosynangiosis (EMS) is a neurosurgical procedure that achieves angiogenesis with low morbidity in patients with moyamoya disease, reducing risk of stroke. However, EMS, surgery has never been studied as an therapeutic option after ischemic stroke. Here we described a novel procedure and feasibility data for EMS after ischemic stroke in mice.

**Methods:** A 60 mins of middle cerebral artery occlusion (MCAo) was used to induce ischemic stroke in mice. After 3-4 hours of MCAo onset/sham, EMS was performed. Mortality of EMS, MCAo and. MCAo+EMS mice was recorded up to 21 days after surgery. Graft tissue viability was measured using a nicotinamide adenine dinucleotide reduced tetrazolium reductase assay.

**Results:** EMS surgery after ischemic stroke does not increase mortality compared to stroke alone. Graft muscle tissue remained viable 21 days after surgery.

**Conclusions:** This novel protocol is effective and well-tolerated, may serve as novel platform for new angiogenesis and thus recovery after ischemic stroke. If successful in mice, EMS can a very feasible and novel treatment option for ischemic stroke in humans.

## Introduction

Ischemic stroke is an acute neurovascular injury with devastating chronic sequelae. Most of the 650,000 yearly stroke survivors in the US are left with permanent functional disability^1^. No available treatments confer neuroprotection and functional recovery after ischemic stroke. Enhancing angiogenesis is a promising therapeutic target, however, previously-studied methods for promoting post-stroke angiogenesis have been limited by unacceptable levels of toxicity or translatability^2^.

Encephalomyosynangiosis (EMS) is a surgical procedure that enhances angiogenesis in moyamoya disease, a condition of narrowed cranial arteries that often leads to stroke. EMS involves partial detachment of a vascular section of the patient’s temporalis muscle from the skull, followed by craniotomy and grafting of the muscle onto the affected cortex. This procedure is well-tolerated and induces cerebral angiogenesis, reducing the risk of ischemic stroke in these patients^3,4^.

EMS has been studied in chronic cerebral hypoperfusion models in rats^5^. However, EMS using a temporalis muscle graft has never been studied in ischemic stroke in rodents. We have standardized a novel protocol of EMS in mice after ischemic stroke via the middle cerebral artery occlusion model (MCAo). This manuscript serves as a description of methods and proof of concept for the use of this novel approach to EMS in mice after MCAo.

## Material and methods

Supporting data are available from corresponding author.

### Animals

We used 8-to 12-week-old, age-and weight-matched C57B/6 wild-type mice—almost equal numbers male and female (total 45 mice)—purchased from The Jackson Laboratory. Mice were fed standard chow diet and water ad libitum. Standard housing conditions were maintained at constant temperature with a 12-hour light/dark cycle. All experiments were approved by the Institutional Animal Care and Use Committee (IACUC) of University of Connecticut Health and conducted in accordance with US guidelines. A total of 4 mice out of 20 from MCAO only and 8 mice out of 34 from EMS +MCAO group was excluded due to mortality.

### MCAo Procedure

MCAo is a well-characterized model of ischemic stroke in rodents, as described by us and others^6-8^.

### EMS Procedure

#### Animal Characteristics

The following protocol should work in any species or strain of rodent.

#### Equipment list

*please see supplementary file*

#### Pre-operative preparation

1. Autoclave instruments prior to surgery. Sanitize operating surface with 70% ethanol.
2. Warm operating surface to 37°C with electric heating pad.
3. Anesthetize mouse with 4-5% isoflurane for induction and 1.5-2.0% isoflurane for maintenance until end of surgery.
4. Place mouse on its left side on the operating surface.
5. Apply eye ointment to protect the right eye.
6. Shave hair over the surgical field (right lateral cranium between eye and ear) with an electric razor.
7. Clean surgical field with 70% ethanol.

#### Surgery (Figure 1)

1. EMS is performed 3-4 hours after MCAo or sham surgery (EMS-only group).
2. 60 mins after MCAo, mice are randomized to MCAo or MCAo+EMS groups.
3. For MCAo+EMS and EMS-only groups, a 10-15 mm skin incision is made with scissors, extending from 1-2 mm rostral to the right ear to 1-2 mm caudal to the right eye.
4. Skin flaps are retracted. Temporalis muscle and skull should be identifiable.
5. Temporalis muscle is bluntly dissected away from the skull. A 2-3 mm myotomy directed ventrally along the caudal border of the muscle may be performed to facilitate ventral reflection.
6. A craniotomy 4-5 mm in diameter is performed at the skull underneath the reflected temporalis muscle using a micro drill.
7. Dura mater is removed with tweezers to expose the pial surface of the brain. Extreme caution is used to avoid accidental injury to the brain.
8. The dorsal border of the temporalis muscle is sutured to the subcutaneous tissue of the dorsal skin flap with 6-0 monocryl filaments, making it flush to the exposed cerebral cortex.
9. The skin incision is closed and cleansed with 70% ethanol then povidone iodine.
10. The mouse is placed back into its cage and monitored until recovery from anesthesia, then is returned to its housing facility.

**Figure 1.**
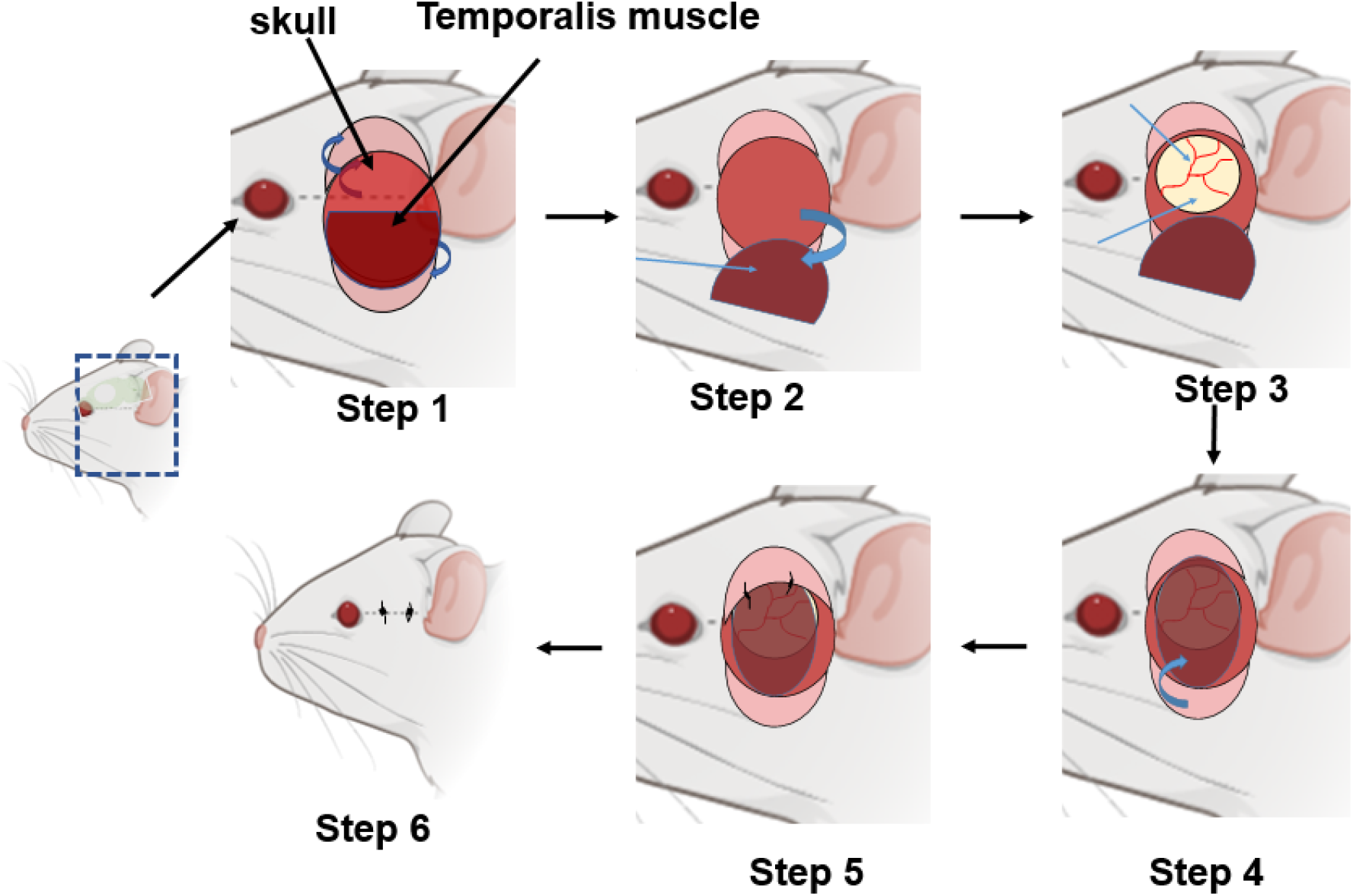
Stepwise EMS after MCAo: 1) Skin incision is made over the right middle cerebral artery territory. Skin and subcutaneous tissues are reflected, exposing skull and temporalis muscle. 2) Temporalis muscle is dissected away from the skull and reflected ventrally. 3) Craniotomy is performed (4-5 mm). Dura is gently removed. 4) Temporalis muscle is placed directly on brain surface to cover exposed cortex. 5) The dorsal edge of the temporalis muscle is sutured to subcutaneous tissue of the dorsal skin flap, flush with the brain surface. 6) The incision is closed, and the mouse is removed from anesthesia and returned to its cage.

#### Post-operative considerations

- Monitor for illness, hydration, and surgical site infection.
- Subcutaneous normal saline (1-2 ml/day) may be prudent.
- Injections, physiologic monitoring, and other testing can proceed without special considerations.
- We follow institutional IACUC guidelines for post-operative analgesia.

#### Graft viability

To assess temporalis muscle graft viability after EMS, a nicotinamide adenine dinucleotide (reduced) tetrazolium reductase (NADH-TR) test was performed as described^9^ with modifications (see Supplementary file). Viability of muscle fibers was calculated as ratio of total area of positive staining to total area of muscle fibers in each picture. Temporalis muscle tissue from the contralateral side was the control.

#### Statistics

Data are presented as mean ± SD. Data were evaluated by student’s t-test (two-tailed unpaired to compare two groups) using GraphPad Prism Software Inc., San Diego, CA).

## Results

### Graft viability and bonding to cortex

Temporalis muscle grafts showed mild but non-significant damage 7 days after surgery (Fig 2), and no difference in viability 21 days after surgery. Using double-immunostaining of anti-α skeletal muscle actin and Lectin-Dy594 (blood vessel marker) antibody, we further showed that temporalis muscle grafts form loose bonds with brain cortex 21 days after EMS (Supplementary Fig 1).

**Figure 2.**
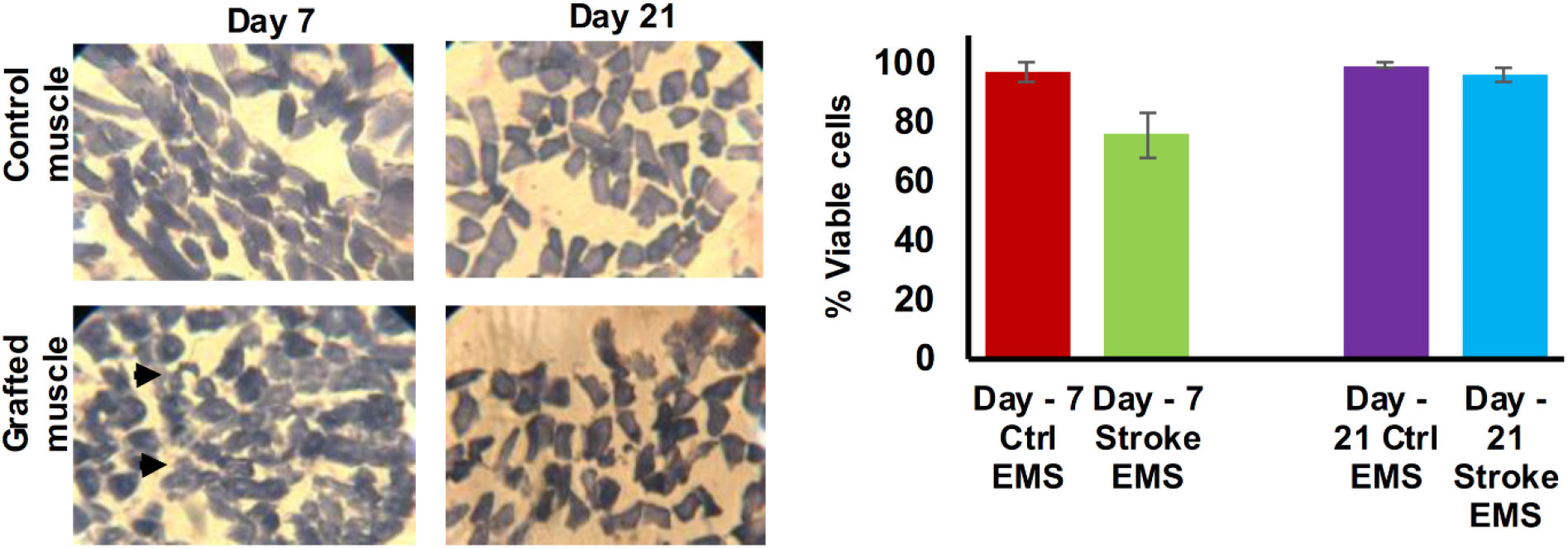
Temporalis muscle grafts maintain viability. (Left) NADH-TR-stained muscle tissue cells from control and grafted muscle 7 and 21 days after MCAo stroke + EMS surgery. Black arrows show damaged cells. (Right) Muscle cells at 7 days after EMS show mild damage (15%, non-significant; t-test) that is no longer present at 21 days. (n=3 mice/time point).

### Mortality outcomes for EMS after stroke

MCAo is an invasive surgical technique that may increase mortality. Here we found 13% mortality in mice 21 days after MCAo surgery, which is an accepted death rate for mice subjected to 60 minutes of MCAo^10^. Performing EMS on mice after MCAo did not significantly increase mortality (16% vs. 13%) (Fig 3).

**Figure 3.**
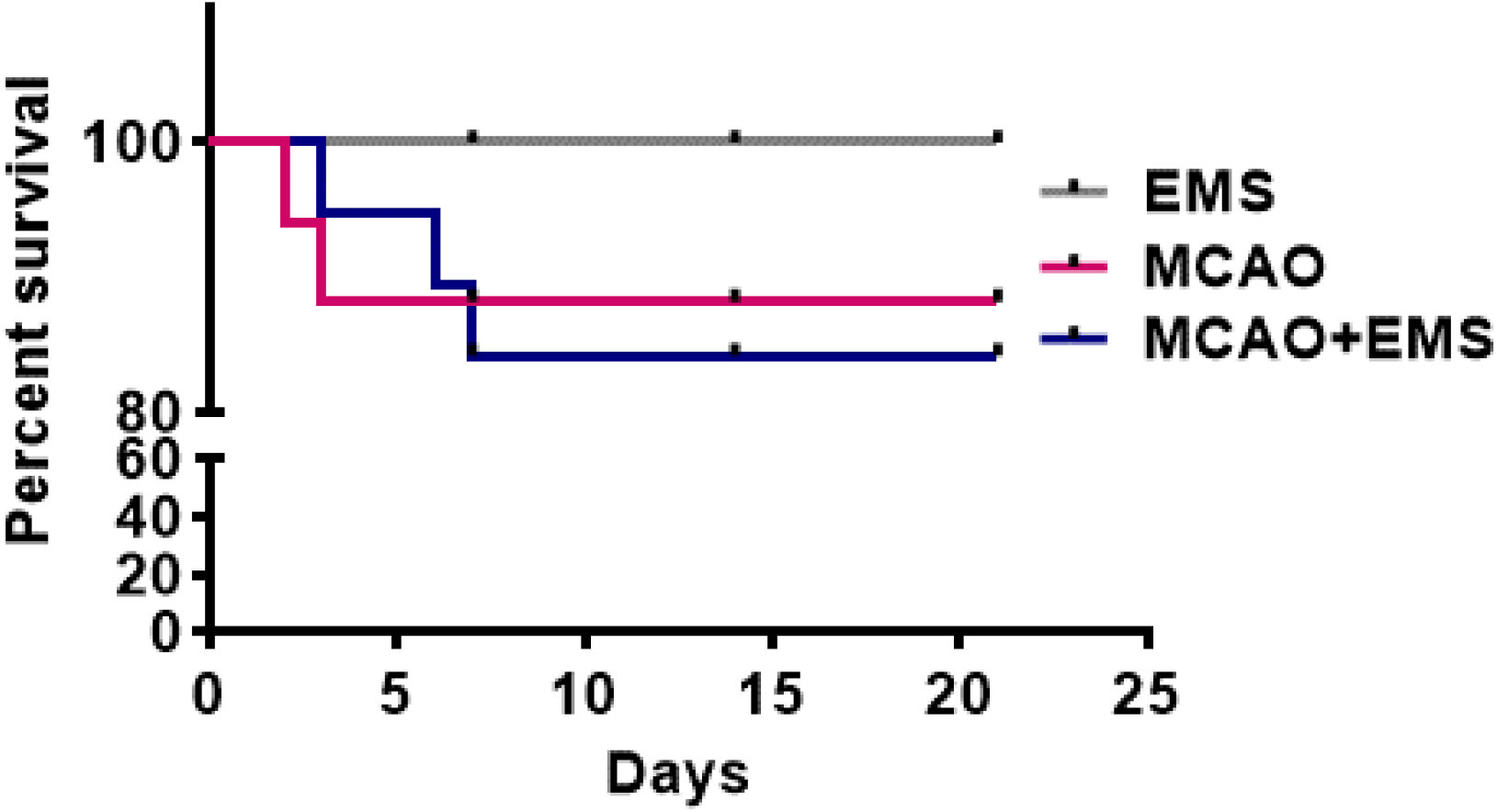
EMS did not increase mortality after stroke (MCAo). Kaplan Meier survival curve shows that EMS+MCAO did not significantly increase post-stroke mortality vs. MCAO alone (16% in EMS+MCAO vs 13% in MCAO). EMS (n=6); MCAO (n=16); MCAO+EMS (n=26).

## Discussion

Here we performed a successful EMS procedure in a mouse model of MCAo-induced stroke. We showed that grafted tissue remains viable and can form loose bonds with brain cortex tissue long after EMS surgery. These findings support the rationale for using cerebral muscle grafts to gradually develop a richly vascular, trophic environment at the site of stroke and is promising for potentially repairing infarcted cerebral tissue in the same environment. Mortality data show that the EMS procedure is well-tolerated by mice even in the physiologically fragile state of recent ischemic stroke.

Our EMS model offers a safe method of achieving cerebral angiogenesis for preclinical study, obviating the need for pharmacologic interventions, which often lead to unwanted side effects or uncontrolled angiogenesis. Further, EMS may provide self-balanced and optimum pro-and anti-angiogenic nutritive factors released by the autologous temporalis muscle tissue, which may further provide support for survival of new blood vessels and new neurons after stroke. EMS has established use in humans and relies on only autologous muscle tissue, increasing the translatability of this angiogenesis model compared to previously-studied xenografts, gene therapy techniques, and intracerebral injections. Many patients with large ischemic strokes require a hemicraniectomy as part of the regimen to manage increasing intracranial pressure. This EMS procedure, which also includes hemi-craniectomy in mice for muscle grafting, may provide preclinical proof of concept for translational application of EMS. Therefore, this model has the potential to expand the knowledge of neurovascular recovery after ischemic stroke as well as to facilitate the development of urgently needed innovation in therapeutics for stroke survivors.

