## supplementary files for "A Novel Model for Encephalomyosynangiosis Surgery after Middle Cerebral Artery Occlusion-Induced Stroke in Mice"

**Equipment required for surgery:**

- Operating microscope
- Operating surface
- Electric heating pad for operating surface
- Micro drill (Harvard Apparatus)
- Isoflurane delivery apparatus
- Isoflurane anesthesia
- Surgical scissors
- Microdissecting tweezers, curved x2
- 6-0 monocryl suture
- Needle driver
- Clamps for tissue retraction
- Rectal thermometer
- 70% ethanol to sanitize operating surface
- Saline or 70% ethanol for irrigation
- Povidone iodine solution
- Ointment for eye protection
- Small electric razor to shave operative site

**Cryosectioning and nicotinamide adenine dinucleotide (reduced)-tetrazolium reductase (NADH-TR) staining protocol:**

At the time of sacrifice, the grafted muscle flap was carefully excised, fixed, and cryopreserved. Several 12 μm-thick cryosections of temporalis muscle tissue were stained for NADH-TR enzyme-histochemical reaction. Slides were incubated for 30 minutes at 37°C in a solution of nitroblue tetrazolium (1.8 mg/dL) and NADH (15 mg/dL) (Sigma-Aldrich Inc, St. Louis, MO) in 0.05 M TRIS buffer (pH 7.6). Unused tetrazolium reagent was removed using increasing, followed by decreasing, concentrations of acetone. Quantitative assessment of NADH-tetrazolium-stained muscle was performed on images from 3 different cross sections and 5 different random fields in each section at 400× magnification.

**Immunohistochemistry:**

Immunostaining was used to visualize the bonding between the brain cortex and grafted skeletal muscle. Additional cryosectioned brain slices (12 μm) were mounted on glass sides, fixed briefly with 4% paraformaldehyde, and washed. Antigen retrieval was done using citrate buffer (pH 6.0), and sections were incubated with blocking buffer followed by incubation overnight with primary antibodies [anti-alpha skeletal muscle actin (green) 1:200; and Lectin-Dy594 (red) antibody]. Three coronal brain sections per mouse (n = 3) were taken 0.40 and 0.90 mm from bregma, stained, and visualized for qualitative analysis. DAPI staining was used to determine the number of nuclei and to assess gross cell morphology.

**Muscle grafts make loose bonds with brain tissue:**

Successful grafting of the temporalis muscle onto the brain cortex is a foremost requirement for the success of this model. Using the EMS-only model, we observed temporalis muscle grafts adhering to cortical surfaces of EMS-only mice (Supplementary Fig 1) suggesting successful EMS surgery, graft implantation, and bonding.

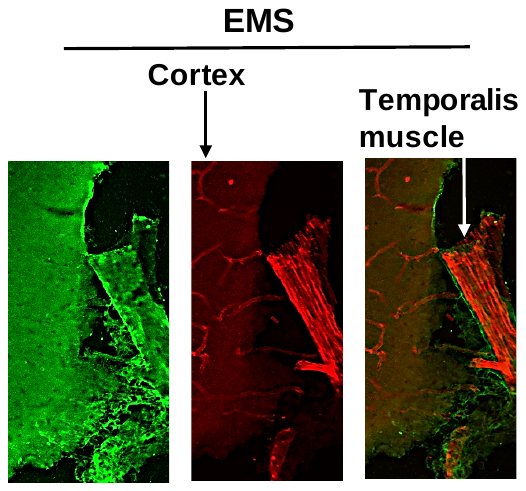

**Supplementary Figure 1**: Bonding of grafted temporalis muscle with brain cortex after EMS-only surgery. EMS tissues stained with anti-alpha skeletal muscle actin (green) and Lectin-Dy594 (red; blood vessel marker) antibody.
